# PTHrP buffers Wnt/β-catenin activity through a negative feedback loop to maintain articular cartilage homeostasis

**DOI:** 10.1101/2022.11.25.517940

**Authors:** Wenxue Tong, Jiankun Xu, Qiuli Qi, Hongjiang Chen, Tao Huang, Chunxia Chen, Weiyang Liu, Zhonglian Huang, Youbin Chen, Zebin Ma, Di Zhao, Jun Hu

## Abstract

Osteoarthritis (OA) is the most common joint disease worldwide and a leading cause of disability. The Wnt/β-catenin cascade is essential in articular cartilage development and homeostasis. It has proved that both overexpression and loss of β-catenin lead to cartilage degeneration and OA symptoms. However, the mechanism of Wnt/β-catenin balance in healthy cartilage remains unclear. In the present work, we confirmed that the Wnt/β-catenin activation and PTHrP suppression in cartilage during the post-traumatic OA process. Then, we demonstrated that Wnt/β-catenin upregulated PTHrP expression through binding to its promoter (P2), and induce mRNA (AT6) transcript expression, while PTHrP repressed Wnt/β-catenin activity, and formed a Wnt/β-catenin-PTHrP negative feedback loop in the very primary chondrocytes to maintain cartilage homeostasis. However, this negative feedback loop vanished in dedifferentiated chondrocytes, hypertrophic chondrocytes, and IL-1β treated very primary chondrocytes. We further found that miR-106b-5p was increased in these “aberrant” chondrocytes and directly targeted PTHrP mRNA to abolish the feedback loop. PKC-ζ was activated by PTHrP through phosphorylation at Thr410/403, and subsequently induced β-catenin phosphorylation and ubiquitination. Finally, we disclosed that exogenous PTHrP attenuated OA progression exogenous PTHrP attenuated OA progression. Together, these findings reveal that PTHrP is a vital mediator to keep Wnt/β-catenin activity homeostasis in healthy cartilage through a negative feedback loop, and PTHrP might be a therapeutic target for OA and cartilage regeneration.

## Introduction

Osteoarthritis (OA) is the most common form of arthritis and affects over 300 million people worldwide (*1, 2*). OA is characterized by the degeneration of articular cartilage, intra-articular inflammation, alterations of periarticular and subchondral bone, and osteophyte formation. The socioeconomic impact of OA and its prevalence is increasing because the population is aging, and obesity is more prevalent in our population. However, the current treatment strategies for OA are still limited to weight loss, physiotherapy, regular exercise, usage of painkillers and anti-inflammatory drugs to relieve the symptoms (*3-5*). Wnt signaling pathway is involved in many biological processes from early embryonic development to organogenesis, to postnatal tissue homeostasis, and plenty of diseases (*6-8*), including musculoskeletal disorders such as osteoporosis (*9*), OA (*10*), and fracture healing (*11*). The Wnt pathway is classified into the canonical Wnt/β-catenin pathway and two noncanonical pathways (*12*). The role of Wnt/β-catenin pathway in articular cartilage during OA process is still contradictory. Cartilage-specific activation of β-catenin resulted in joint destruction together with the loss of stable chondrocyte phenotype and showed terminal hypertrophic morphology that was associated with OA-like lesions (*13*). In a mouse model, the activation of β-catenin in cartilage induced proteoglycan loss while abnormally increased cartilage thickness and chondrocyte proliferation rate, and ablation of β-catenin in cartilage led to hypocellularity (*14*). Furthermore, the disfunction of β-catenin in cartilage led to chondrocyte apoptosis and OA-like symptom (*15*). **Thus, both overexpression and loss of** β**-catenin lead to cartilage degeneration and OA symptoms**. Briefly, the gain of β-catenin function leads to chondrocyte hypertrophy and reduction in matrix quality, while the loss of β-catenin function leads to chondrocyte death and tissue damage (*12*). Some certain Wnt ligands, such as Wnt16 and Wnt5a, were considered as the buffer factors for Wnt/β-catenin activity in articular chondrocytes because they can both activate and suppress Wnt/β-catenin pathway (*16, 17*), and this also indicates the crosstalk between canonical and non-canonical Wnt pathways (*12*). Several inner factors, like Axin1 and Axin2, also form a general negative feedback loop for the canonical Wnt/β-catenin pathway (*18*). **However, a complete regulatory system and mechanism for the homeostasis of Wnt/**β**-catenin that specifically in health articular cartilage remains unclear**.

Parathyroid hormone-related protein (PTHrP) and Parathyroid Hormone (PTH) share limited sequence homology, and they play a similar biological function through the same receptor, parathyroid hormone 1 receptor (PTH1R). PTH is mainly expressed and secreted by parathyroid master cells, while PTHrP can be found in the thyroid, parathyroid gland, bone marrow, cardiovascular, brain, skin, bone, and other tissues. Unlike PTH, which regulates the balance of bone metabolism throughout the whole body via the circulatory system, PTHrP mainly regulates the biological activity of adjacent tissues in the form of autocrine or paracrine (*19*). PTHrP plays an essential role in endochondral ossification during long bone development through a typical Ihh-PTHrP negative feedback loop. The delicate balance between Ihh and PTHrP tightly controls the endochondral ossification process: Ihh promotes chondrocyte hypertrophy, while PTHrP, by contrast, inhibits chondrocyte hypertrophy; meanwhile, Ihh induces the expression of PTHrP in chondrocytes through Gli1/2. Thus, the activity of Ihh on chondrocyte hypertrophy is delicately balanced by PTHrP during the endochondral ossification process, especially in the secondary ossification center and the postnatal epiphyseal plate (*20*). The chondrogenic transcription factor Sox9 is a target of PTHrP in the epiphyseal plate of endochondral bones (*21*). Previous studies, including ours (*22*), have shown that PTHrP is mainly expressed in the superficial and middle layers of articular cartilage, while it is barely detected in the deep and calcified layers (*23-25*). Both PTHrP (*22*) and its receptor, PTH1R (*26*), in cartilage are dramatically downregulated during OA progression. The mice of PTHrP knockout in cartilage showed more severe cartilage degradation in the post-traumatic OA model than in the wild-type mice (*23*). PTHrP in articular cartilage was about 5 times higher than that in osteophytes, suggesting that PTHrP is an important factor for cartilage homeostasis, while insufficient PTHrP lead to chondrocyte hypertrophy and endochondral ossification induced osteophyte formation (*27*). Furthermore, PTHrP overexpression represses Collagen Type X (Col10), a marker of chondrocyte hypertrophy (*28*). PTH (1-34), an amino-terminal fragment of PTH that has a similar function to PTHrP, achieved the same cartilage protection effect as PTHrP (*29*). These studies disclosed the role of PTHrP for cartilage protection (*30*).

The crosstalk between the noncanonical Wnt/PCP pathway and PTHrP in articular cartilage has been disclosed in our previous study that Wnt/PCP induced PTHrP expression through mTORC1 to attenuate OA (*22*). Meanwhile, the interaction of canonical Wnt/β-catenin and PTHrP during endochondral ossification of skeleton development has been revealed that Wnt/β-catenin inhibited PTHrP activity, but did not affect its expression, to initiate chondrocyte hypertrophy (*31*). However, the mutual promotion effect between Wnt/β-catenin signaling and PTHrP has also been demonstrated (*32, 33*). Here, we investigated the mechanism for Wnt/β-catenin balance in healthy cartilage and found that PTHrP is a crucial mediator to form a negative feedback loop that maintains the Wnt/β-catenin activity homeostasis. This study revealed that PTHrP is a potential therapeutic target for OA, and cartilage protection and regeneration.

## Results

### Wnt/β-catenin was over-activated, while PTHrP was suppressed in OA cartilage

We first detected the expression of β-catenin and PTHrP in the destabilization of the medial meniscus (DMM) induced mouse OA samples and clinical human cartilage specimens. DMM surgery induces dramatic OA progression compared to the sham control group, as shown by cartilage morphology (Fig. 1A) and OARSI grade and score (Fig. 1B). The clinical cartilage specimens of the loading area, medial femoral condyle (MFC), showed cartilage degeneration. In contrast, the non-loading area, lateral femoral condyle (LFC), still has the intact cartilage morphology (Fig. 1C). The expression of β-catenin was significantly upregulated in the DMM OA cartilage and MFC cartilage, compared to the sham control group and LFC cartilage, respectively (Fig. 1D-E). However, PTHrP showed a reversed variation trend as that of β-catenin, and significantly reduced in the DMM OA cartilage and MFC cartilage (Fig. 1D-E). These data indicated a potential linkage between β-catenin and PTHrP.

**Figure 1.**
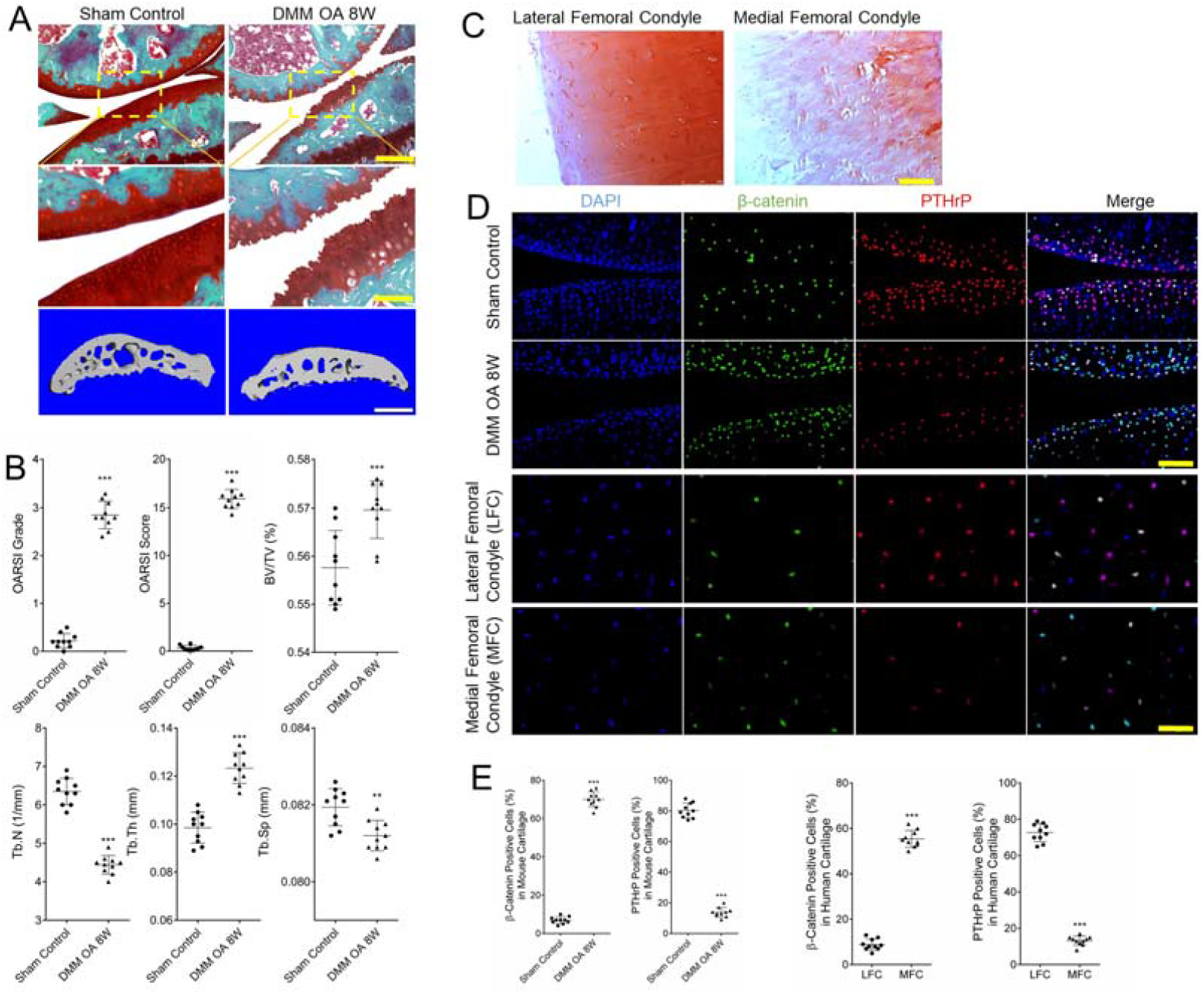
Wnt/β-catenin was over-activated, while PTHrP was suppressed in OA cartilage. (A) Safranin O and fast green staining of representative paraffin sections and micro-CT assay of the mouse knee samples and (B) OA evaluation by OARSI grade and score, and micro-CT score showed the significant OA progression after DMM surgery. Scale bar, 250 μm (top, yellow), 80 μm (bottom, yellow). (C) The morphology of lateral femoral condyle (LFC) cartilage and medial femoral condyle (MFC) cartilage in specimens from total knee replacement surgery patients. Scale bar, 250 μm. (D, E) The expression of β-catenin was significantly upregulated in the DMM OA cartilage and MFC cartilage, compared to the sham control group and LFC cartilage. Scale bar, 50 μm. \*\*\**p* <0.001.

### Wnt/β-catenin upregulated PTHrP expression, while PTHrP repressed Wnt/β-catenin activity in chondrocytes

To further examine the connection of β-catenin and PTHrP, we isolated clinical human articular chondrocytes and used the very primary chondrocytes (P0) that showed typical cobblestone morphology for experimental assay. The very primary chondrocytes were treated with Wnt3a, a classic ligand for the Wnt/β-catenin pathway, and the PTHrP expression was significantly upregulated both in mRNA and protein level. The targeting genes, Nkx3.2 and Zfp521, were also upregulated by Wnt3a (Fig. 2A-B). However, when we detected the expression of Wnt/β-catenin targeting genes, their expression was not sustainedly upregulated by high dosage (20, 40, 80 ng/ml) of Wnt3a (Fig. 2C-D) but maintained on a stable high level. This phenomenon indicated a buffer system might exist to keep Wnt/β-catenin activity. To examine if the PTHrP is responsible for this phenomenon, we interfered PTHrP expression by PTHrP-siRNA. The results showed that PTHrP-siRNA significantly reduced the PTHrP expression and its targeting genes (Fig. 2E-F). Three distinct PTHrP proteins PTHrP 1-139, PTHrP 1-173, and PTHrP 1-141 was encoded ty three alternative *PTHrP* transcripts AT5.2, AT6, and AT7 (Fig. 2G). We then determined which of the three transcripts was induced by Wnt3a in the very primary chondrocytes via transcript-specific QRT-PCR assay. The results showed that Wnt3a treatment of human very primary chondrocytes selectively induced the AT6 transcript encoding PTHrP 1-173 instead of AT5.2 or AT7 transcripts (Fig. 2H). The β-catenin-ChIP analyses of *PTHrP* gene promoters revealed that β-catenin bound to the P2 promoter region, but not to the P1 and P3 promoter regions (Fig. 2I). The Wnt/β-catenin activity was sustainedly upregulated by a high dosage of Wnt3a together with PTHrP-siRNA showed by QRT-PCR (Fig. 2J), western blotting (Fig. 2K), and immunofluorescence (Fig. 2L). These results suggested that Wnt/β-catenin directly upregulated PTHrP expression, while PTHrP repressed Wnt/β-catenin activity, and PTHrP was a critical mediator to buffer Wnt/β-catenin activity in the very primary chondrocytes.

**Figure 2.**
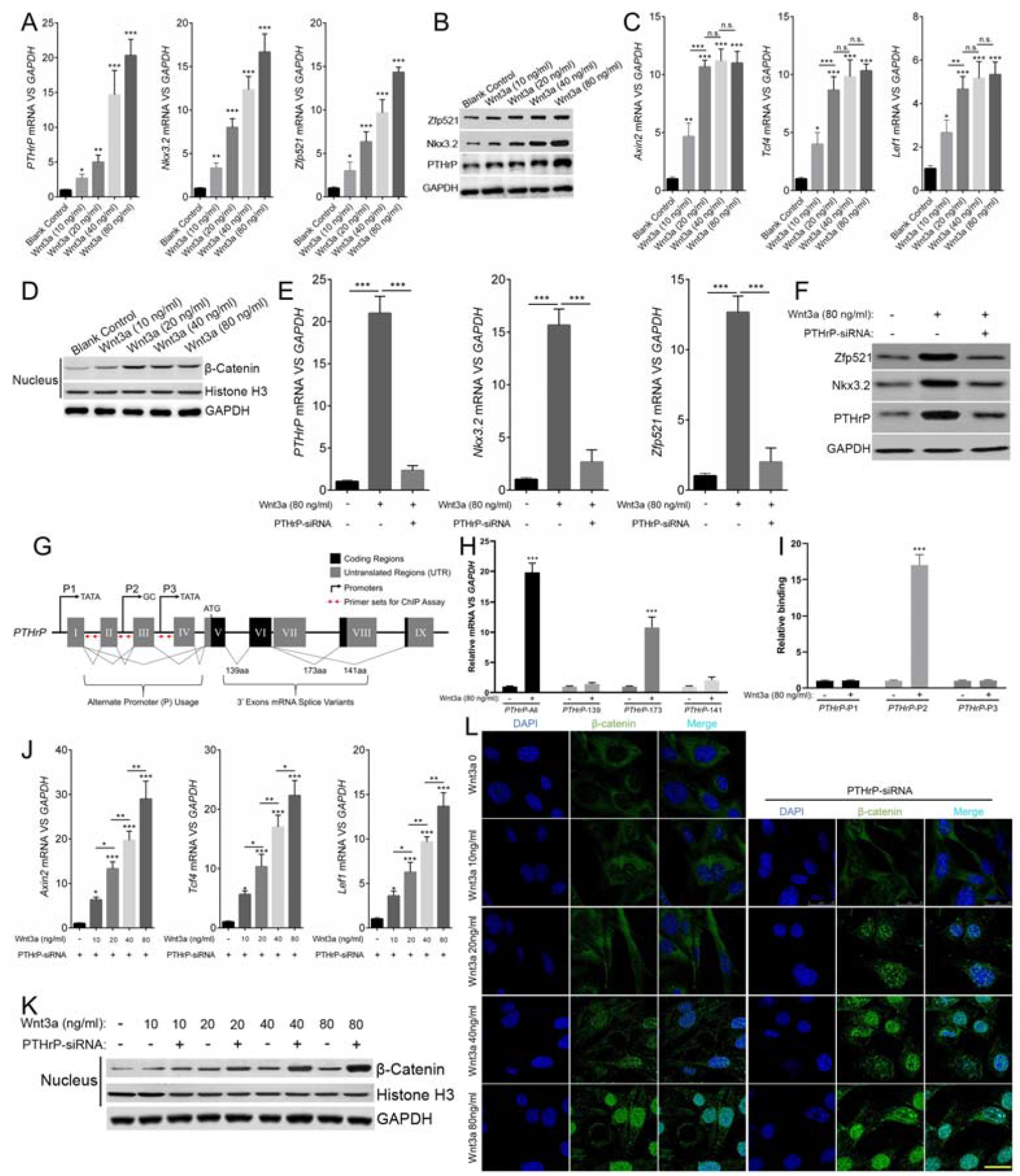
Wnt/β-catenin upregulated PTHrP expression, while PTHrP repressed Wnt/β-catenin activity in chondrocytes. (A, B) PTHrP and its targeting genes expression were upregulated by Wnt3a. (C, D) Wnt/β-catenin activity was increased by Wnt3a, but not sustainedly upregulated by high dosage (20, 40, 80 ng/ml) of Wnt3a. (E, F) PTHrP-siRNA significantly reduced the PTHrP expression and its targeting genes. (G) Three alternative promoters and nine exons are included in the *PTHrP* gene. Alternative splicing of 5’-exons produce transcripts AT5.2, AT6, or AT7 (*SpliceSeq* nomenclature) encoding PTHrP 1-139, 1-173, or 1-141 protein as showed by the lines and black boxes, respectively. The three promoters are located upstream of exon I (P1), between II and III (P2), and between III and IV (P3). (H) QRT-PCR analysis using primers specific to transcripts AT5.2, AT6, or AT7 indicated that Wnt3a preferentially induced AT6 encoding PTHrP 1-173 in the human very primary chondrocytes. (I) ChIP assay of P1, P2, or P3 sequences captured by anti-β-catenin antibody or control IgG on sheared chromatin from Wnt3a stimulated human very primary chondrocytes showed inducible enrichment of P2 promoter. (J-L) Wnt/β-catenin activity was sustainedly upregulated by high dosage of Wnt3a together with PTHrP-siRNA. \**p* < 0.05, \*\**p* < 0.01, \*\*\**p* < 0.001. Scale bar, 25 μm.

### The Wnt/β-catenin-PTHrP negative feedback loop was not detected in dedifferentiated chondrocytes, hypertrophic chondrocytes, and IL-1β treated very primary chondrocytes

To examine if the buffer effect of PTHrP for Wnt/β-catenin is a common role, we repeated relevant experiments in the dedifferentiated fibrous chondrocytes (DFC), hypertrophic chondrocytes (HC), and IL-1β treated very primary chondrocytes (IL-1β). The PTHrP expression and the downstream targeting genes were significantly decreased in all these three groups compared with the very primary chondrocytes (PC) group (Fig. 3A-B). In these situations, the Wnt/β-catenin activity was increased to a dramatic level compared to the PC group. However, when the PTHrP cytokine was added, the Wnt/β-catenin activity was significantly suppressed (Fig. 3C-E). Moreover, the dedifferentiated fibrous chondrocytes were partially rescued by PTHrP in a dose-dependent manner as shown by the chondrocyte markers including Sox9, Collagen 2a1 (Col2a), aggrecan, and Hyaluronic acid synthase 2 (Has2), and the fibrous cartilage marker Collagen 1a1 (Col1a1) (Fig. 3F-G). The upregulated PTHrP and its targeting genes expression by Wnt3a was also significantly eliminated in the DFC, HC, and IL-1β group (Fig. 3H-I, labeled by # to show the statistical difference versus the same treatment of PC). These data indicated that the buffer role of PTHrP disappeared in dedifferentiated fibrous chondrocytes, hypertrophic chondrocytes, and IL-1β treated very primary chondrocytes because its expression was strongly repressed.

**Figure 3.**
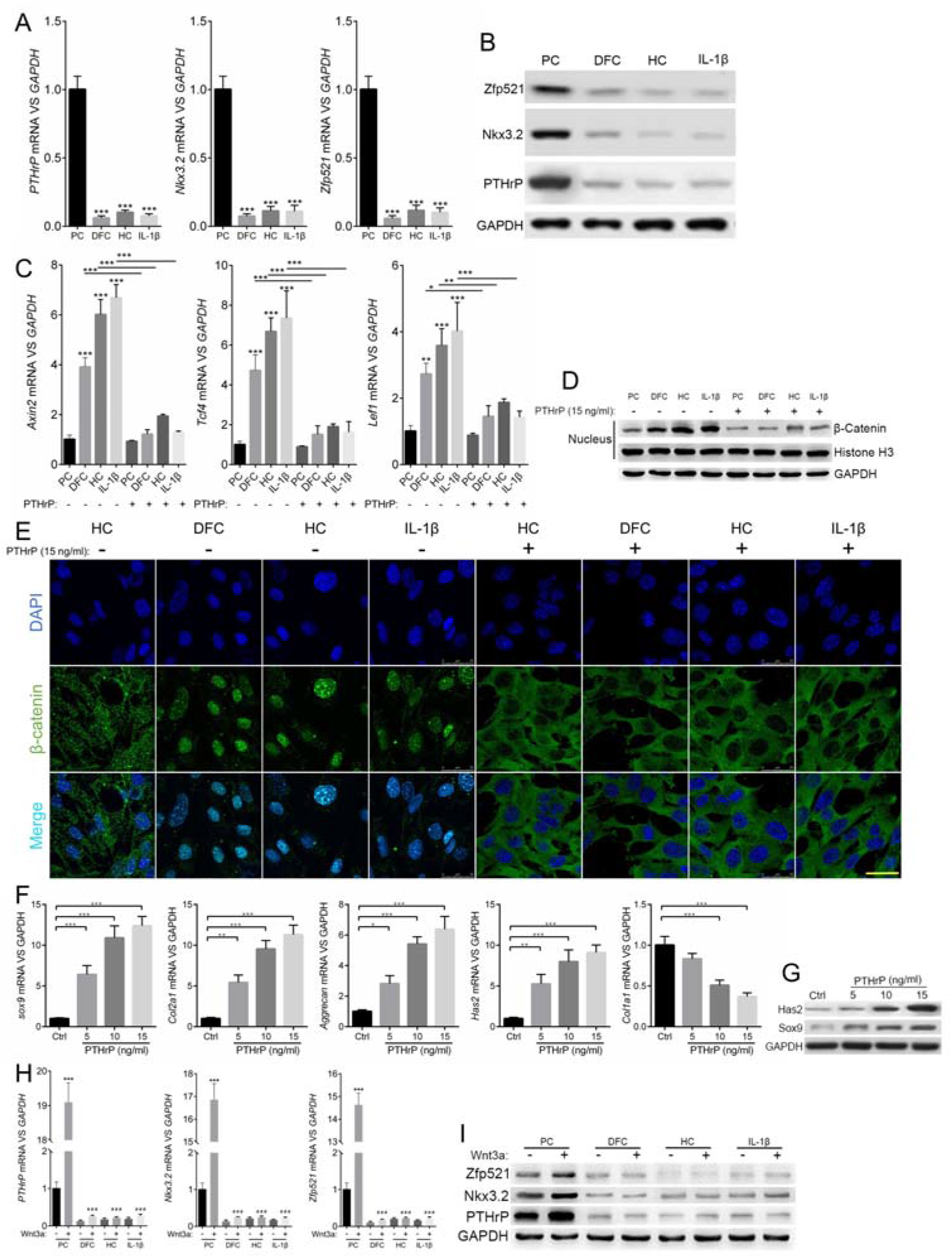
The Wnt/β-catenin-PTHrP negative feedback loop was not detected in dedifferentiated chondrocytes, hypertrophic chondrocytes, and IL-1β treated very primary chondrocytes. (A, B) The PTHrP expression and the downstream targeting genes were significantly decreased in the dedifferentiated fibrous chondrocytes (DFC), hypertrophic chondrocytes (HC), and IL-1β treated very primary chondrocytes (IL-1β) groups compared with the very primary chondrocytes (PC) group. (C-E) Wnt/β-catenin activity was significantly suppressed by PTHrP. (F, G) The dedifferentiated fibrous chondrocytes were partially rescued by PTHrP in a dose dependent manner showed by the chondrocyte markers. (H, I) The DFC, HC and IL-1β treatment eliminated the Wnt3a upregulated PTHrP and its targeting genes expression. \**p* < 0.05, \*\**p* < 0.01, \*\*\**p* < 0.001. (For figure 3H, * means the comparison versus the (Wnt3a: –) in the same cell type group, and # means the comparison versus the (Wnt3a: +) in the PC group). Scale bar, 25 μm.

### The miR-106b-5p was upregulated in the “aberrant” chondrocytes and directly targeted PTHrP mRNA

To explore the reason for PTHrP downregulation, we systemically analyzed a series of micro-RNAs that directly target PTHrP mRNA. 45 micro-RNAs were chosen as the candidates according to *targetscan.org* and *mirdb.org* database and the score rankings. Among these micro-RNAs, the hierarchical cluster analysis showed that the miR-106b-5p was significantly upregulated in all the three “aberrant” chondrocytes (Fig. 4A). Overexpression of miR-106b-5p inhibited PTHrP expression and increased Wnt/β-catenin activity in the very primary chondrocytes (Fig. 4B-C). While in the IL-1β treated very primary chondrocytes, knockdown of miR-106b-5p by its inhibitor increased PTHrP expression and suppressed Wnt/β-catenin activity (Fig. 4D-E). The schematic representation of the PTHrP 3’ UTR indicated the binding site of miR-106b-5p, and the mutant strategy on the binding site was also described (Fig. 4F). The relative luciferase activity showed that ectopic expression of miR-106b-5p led to a significant reduction of luciferase activity in the wild-type PTHrP gene 3’ UTR but not that of the mutant reporter (Fig. 4G). These data indicated that miR-106b-5p was upregulated in dedifferentiated chondrocytes, hypertrophic chondrocytes, and IL-1β treated very primary chondrocytes and directly targeted PTHrP mRNA.

**Figure 4.**
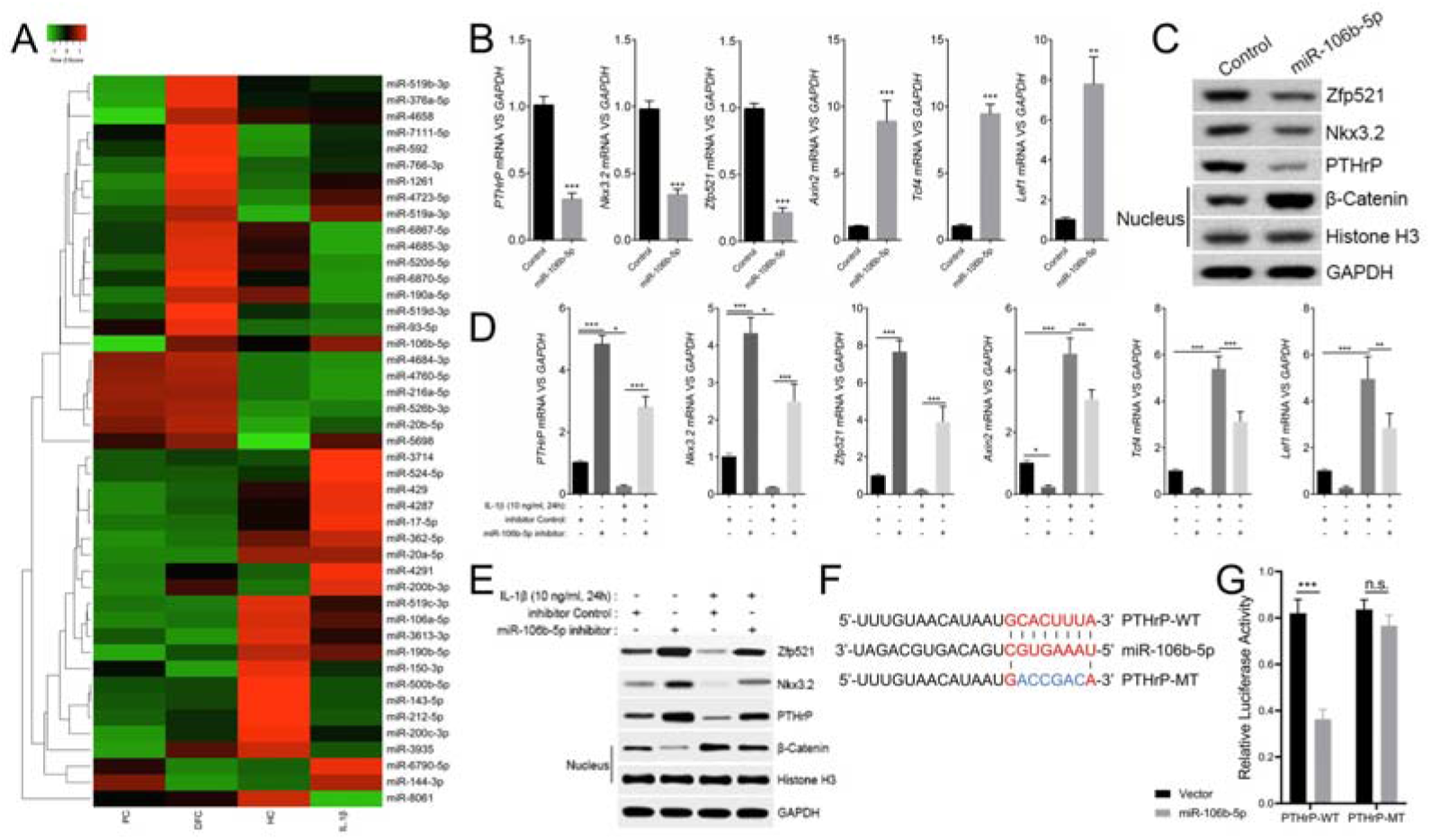
The miR-106b-5p was upregulated in the “aberrant” chondrocytes and directly targeted PTHrP mRNA. (A) The miR-106b-5p was significantly upregulated in DFC, HC and IL-1β treated very primary chondrocytes groups by hierarchical cluster analysis. (B, C) Overexpression of miR-106b-5p inhibited PTHrP expression and increased Wnt/β-catenin activity in the very primary chondrocytes. (D, E) Knockdown of miR-106b-5p by its inhibitor increased PTHrP expression and suppressed Wnt/β-catenin activity in the IL-1β treated very primary chondrocytes. (F) The schematic representation of the PTHrP 3’ UTR binding site of miR-106b-5p, and the mutant strategy. (G) Ectopic expression of miR-106b-5p led to a significant reduction of luciferase activity in the wild type PTHrP gene 3’ UTR but not that of the mutant reporter. \**p* < 0.05, \*\**p* < 0.01, \*\*\**p* < 0.001, n.s. = no significant difference.

### PTHrP activated PKC-ζ to induce β-catenin phosphorylation and ubiquitination

To further explore the mechanism of PTHrP down-regulating Wnt/β-catenin activity, we detected the downstream phosphorylated (p-) PKC isozymes that in response to PTHrP. The p-PKC-ζ (Thr410/403) was increased by PTHrP treatment in the very primary chondrocytes, while the level of p-PKC-α/βII (Thr638/641), p-PKC-δ (Thr505), and p-PKC-ε (Ser729) were not affected by PTHrP (Fig. 5A). While the p-PKC-ζ (Thr410/403) was dramatically decreased in the DFC, HC and IL-1β treated very primary chondrocytes (Fig. 5B). Moreover, PKC-ζ siRNA impaired the PTHrP decreased Wnt/β-catenin activity in the IL-1β treated very primary chondrocytes (Fig. 5C-E). The rescue of cartilage marker expression by PTHrP in the dedifferentiated chondrocytes was also impaired by PKC-ζ siRNA (Fig. 5F-G). Co-immunoprecipitation (Co-ip) analysis showed the interaction of β-catenin and p-PKC-ζ (Fig. 5H). PTHrP phosphorylates β-catenin on N-terminal of Ser33/Ser37 and Ser45 instead of C-terminal of Ser675, while PKC-ζ siRNA represses the phosphorylation (Fig. 5I). Further analysis showed that IL-1β inhibited β-catenin ubiquitination, and PTHrP induced β-catenin ubiquitination while PKC-ζ siRNA repressed PTHrP’s function (Fig. 5J). These data indicated that PKC-ζ was activated PTHrP, and subsequently induced β-catenin phosphorylation and ubiquitination.

**Figure 5.**
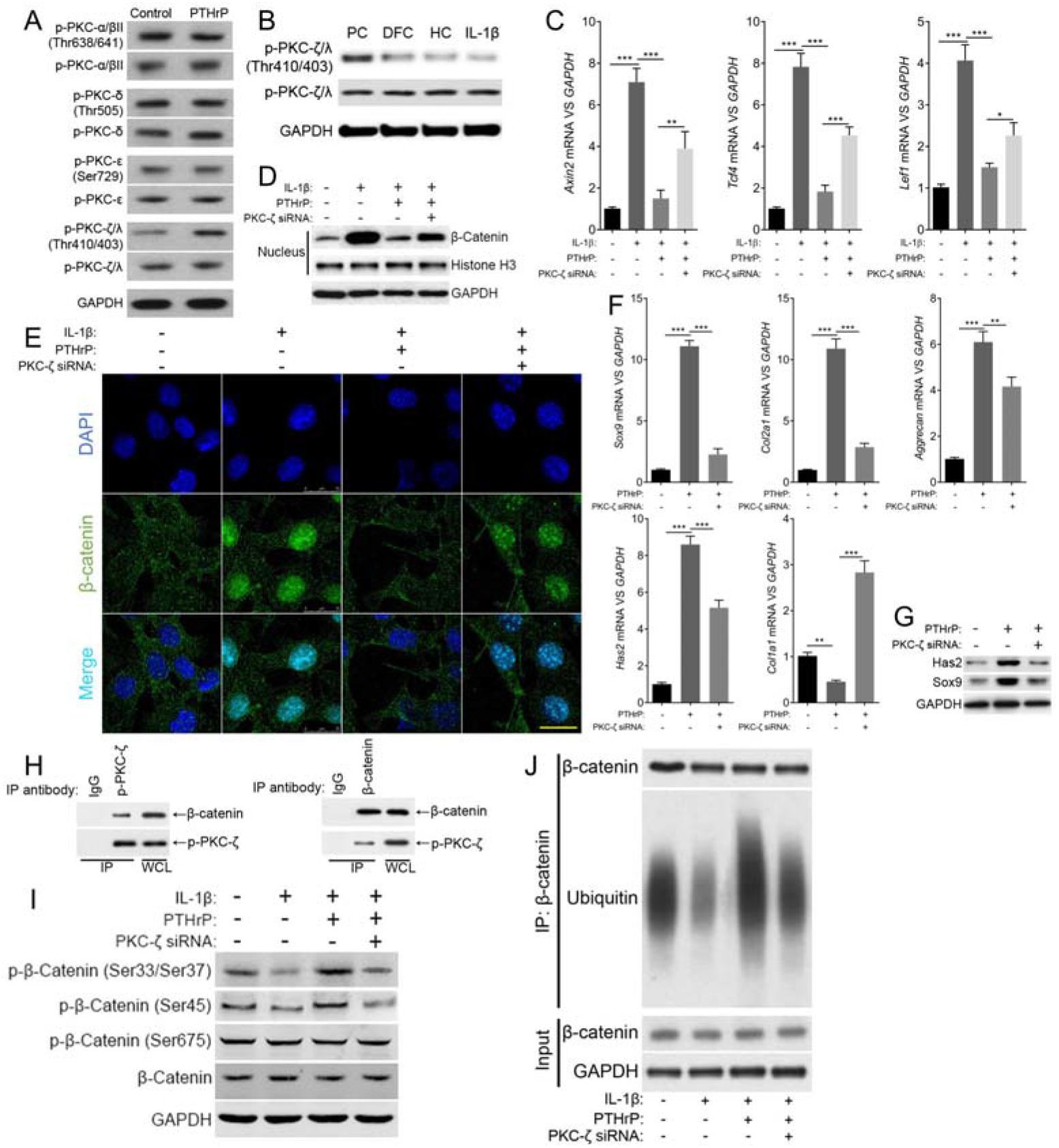
PTHrP activated PKC-ζ to induce β-catenin phosphorylation and repress Wnt/β-catenin activity. (A) The p-PKC-ζ (Thr410/403) was specifically increased by PTHrP treatment in the very primary chondrocytes. (B) The p-PKC-ζ (Thr410/403) was decreased in the DFC, HC and IL-1β treated very primary chondrocytes. (C-E) The PTHrP decreased Wnt/β-catenin activity in the IL-1β treated very primary chondrocytes was impaired by PKC-ζ siRNA. (F, G) PKC-ζ siRNA impaired the rescue of cartilage marker expression in the dedifferentiated chondrocytes by PTHrP. (H) The interaction of β-catenin and p-PKC-ζ was proved by Co-immunoprecipitation (Co-ip). (I) PTHrP phosphorylated β-catenin on N-terminal of Ser33/Ser37 and Ser45 instead of C-terminal of Ser675, while PKC-ζ siRNA represses the phosphorylation. (J) IL-1β inhibited β-catenin ubiquitination, and PTHrP induced β-catenin ubiquitination while PKC-ζ siRNA repressed PTHrP’s function. \**p* < 0.05, \*\**p* < 0.01, \*\*\**p* < 0.001. Scale bar, 25 μm.

### P-PKC-ζ(Thr410/403) was decreased in OA cartilage

To demonstrate that p-PKC-ζ (Thr410/403) is involved in the OA process, we examined its expression in articular cartilage of mouse samples and clinical specimens. The p-PKC-ζ (Thr410/403) expression was significantly reduced in mouse DMM OA cartilage, and in clinical MFC cartilage as compared with that in LFC (Fig. 6A-B). Intra-articular injection of AAV9 delivered PTHrP remarkably attenuated OA progress showed by cartilage morphology (Fig. 6C) and OARSI score (Fig. 6D). The β-catenin was decreased, while p-PKC-ζ (Thr410/403) was increased in articular cartilage by AAV9-*PTHrP* treatment compared with AAV9-*Vector* (Fig. 6E). These data indicated that PTHrP might be a therapeutic target for OA and cartilage regeneration through PKC-ζ activation.

**Figure 6.**
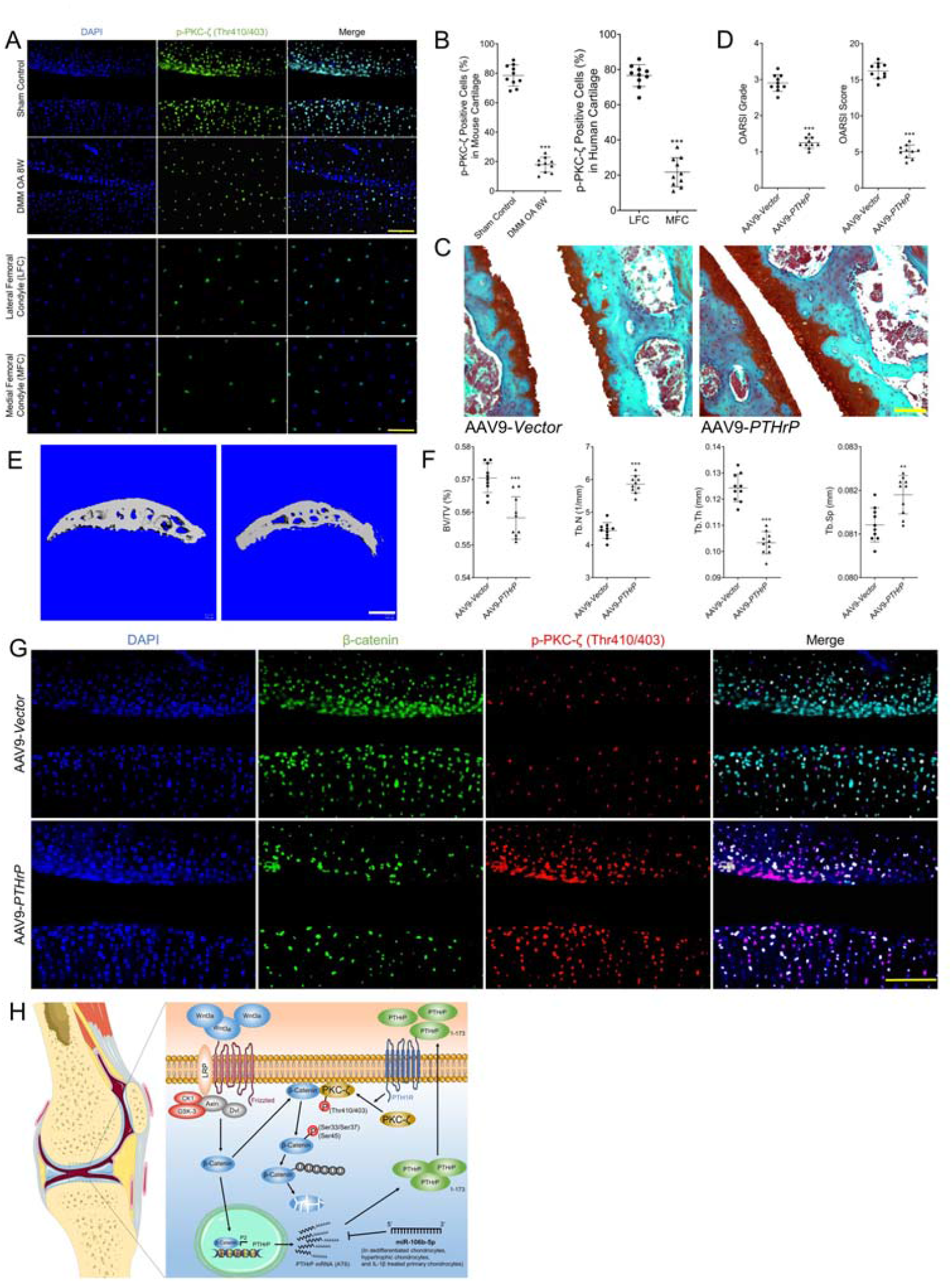
P-PKC-ζ (Thr410/403) was decreased in OA cartilage. (A, B) The expression of p-PKC-ζ (Thr410/403) was significantly upregulated in the DMM OA cartilage and MFC cartilage, compared to the sham control group and LFC cartilage. (C-F) AAV9 delivered PTHrP remarkably attenuated OA progress shown by the cartilage and subchondral bone morphology and score. (G) AAV9-PTHrP decreased β-catenin and increased p-PKC-ζ (Thr410/403) in articular cartilage. (H) Schematic diagram of the Wnt/β-catenin-PTHrP negative feedback loop in chondrocytes of healthy cartilage. Wnt/β-catenin upregulated PTHrP expression by binding to primer P2, while PTHrP repressed Wnt/β-catenin activity. The miR-106b-5p nullify the Wnt/β-catenin-PTHrP feedback loop by targeting PTHrP mRNA (AT6) in dedifferentiated chondrocytes, hypertrophic chondrocytes, and IL-1β treated chondrocytes. PKC-ζ is the critical mediator for PTHrP repressing Wnt/β-catenin activity by inducing the phosphorylation, ubiquitination, and final degeneration of β-catenin. Scale bar, 50 μm in (A) and (E), 80 μm in (C). ***p <0.001.

## Discussion

Both the role of Wnt/β-catenin and PTHrP in skeleton development has been studied for decades. A stable Wnt/catenin level is crucial for cartilage homeostasis since its overactivation or loss will lead to joint damage and OA like symptoms (*12*). The role of PTHrP in inhibiting chondrocyte hypertrophy, especially the canonical negative feedback loop between PTHrP and Ihh during the endochondral ossification, has been well disclosed (*34, 35*). Based on this well-established theory of the role of PTHrP in skeleton development, it has been found that PTHrP participated in the articular cartilage maintenance as its role in growth plate cartilage (*25*). Several studies, including ours, have disclosed that PTHrP located in the superficial and middle zoon of the healthy articular cartilage (*22-25*). However, the application of PTHrP in OA prevention or treatment and cartilage repair has not been extensively considered. In this study, we found a novel function of PTHrP that buffered Wnt/β-catenin activity and formed a similar Wnt/β-catenin-PTHrP negative feedback loop as the Ihh-PTHrP one during the endochondral ossification (Figure 6F). This conclusion provides another explanation for the protection of articular cartilage by PTHrP.

It is still controversial about the role of Ihh-PTHrP negative feedback loop in postnatal articular cartilage. Some studies defaulted that the feedback loop was sustained in the articular cartilage and maintained the expression of PTHrP (*29, 36*). While other studies found that the mechanical loading, instead of Ihh, maintained the PTHrP expression in articular cartilage (*23, 25*). Gli1 and Gli2 are the two transcription factors that directly promote PTHrP expression. It has been proved that Gli2 directly upregulated PTHrP expression independent of Ihh (*37*). Moreover, Ihh was slightly detected in healthy articular cartilage and upregulated during OA progression (*38, 39*), but Gli1 was strongly detected in murine postnatal articular cartilage (*40*). These studies, together with our current one, indicate that the canonical Ihh-PTHrP negative feedback does not exist in postnatal articular cartilage, and the PTHrP’s expression is maintained by Gli1/Gli2 independent of Ihh. The detailed mechanism, such as the crosstalk between the mechanical loading and Gli1/Gli2 activation worth further investigation.

Although the cartilage protection role of PTHrP is a well-established conclusion by many studies. Our current results indicate that it is a very fragile balance system between PTHrP and Wnt/β-catenin, that only existed in healthy articular cartilage. In other words, the situation of articular cartilage is the result of the combination of internal and external factors, and the PTHrP-Wnt/β-catenin negative feedback loop is just one of the factors. At the same time, the cartilage state also affects this feedback loop. Although this PTHrP-Wnt/β-catenin feedback loop was destroyed during OA progression, as showed that PTHrP was significantly down-regulated while β-catenin was upregulated, it was still partially rescued by exogenous PTHrP supplement, and accordingly, the OA development was also significantly attenuated. These results indicated the potential application of PTHrP as a supplement delivered by an efficient and safe cartilage-targeting vector.

According to our research data, the miR-106b-5p was commonly and significantly upregulated in dedifferentiated fibrous chondrocytes, hypertrophic chondrocytes, and IL-1β treated very primary chondrocytes. The further study disclosed that miR-106b-5p destroyed the PTHrP-Wnt/β-catenin feedback loop by directly targeting PTHrP in chondrocytes. The micro-RNAs family constitutes a huge regulatory system, and hundreds of micro-RNAs will be altered when the cell or environment state changed (*41*). In this study, we performed the systemic analysis and found that miR-106b-5p is the most common altered one among the candidates that targeting PTHrP. However, as the results demonstrated, several micro-RNAs also significantly altered in one or two situations. These micro-RNAs may also play an essential role by repressing PTHrP pathway activity in these certain situations, and their detailed role and regulatory manner merit further studies, respectively.

PKC has been in the limelight since the discovery four decades ago. There are three subfamilies of PKC family according to their domain structure: conventional PKC (cPKC), novel PKC (nPKC), and atypical PKC (aPKC). Ten family members of PKC have been identified (*42*). The PKC proteins undergo complicated variations during chondrogenesis and OA progression (*43-46*). But most studies and reviews focused on the individual subfamilies instead of specific PKC proteins. Thus, the function of each PKC protein in OA is still far more from understanding. In consideration that PTHrP is the target of non-canonical Wnt/planar cell polarity (PCP) pathway in chondrocytes (*22*), and the regulatory function of aPKC in cell polarity (*47, 48*), a much more complicated network among Wnt/β-catenin, PTHrP, Wnt/PCP and aPKC may exist during skeleton development and OA progression. Moreover, given the well-established toolbox for PKC pharmacology especially in the cancer research field (*49*), a refined study of the specific PKC proteins in OA may provide us available drugs that will be potentially applied in OA prevention and treatment. In conclusion, our work suggests a PTHrP-Wnt/β-catenin feedback loop that involved PKC-ζ, and they are the potential therapeutic targets for OA prevention and treatment.

## Materials and Methods

### Clinical Specimens preparation

The clinical osteoarthritis (OA) specimens were collected from 10 total knee joint replacement patients (7 male patients and 3 female patients, 70-78 years old). The specimens were classified into medial femoral condyle (MFC) that in the loading area and lateral femoral condyle (LFC) that aside of the loading area of the knee joint.

### Primary Chondrocytes Isolation and Culture

The isolation and culture of human very primary chondrocytes were performed according to established protocol. The intact LFC area of the articular cartilage was rinsed twice with sterile PBS and cut into small pieces. The pieces were digested in 0.25% trypsin-EDTA for 20 minutes and 0.02% Collagenase II for 5 hours at 37°C in a thermal incubator under 5% CO_2_. The isolated cells were filtrated through a 70 μm diameter filter and seeded onto a culture dish with a DMEM: F12=1:1 medium contained 10% FBS, 1% penicillin-streptomycin.

### Total RNA Extraction

The TRIzol Reagent (1 mL, Thermo Fisher Scientific) was added to the cultured cells and lysis at room temperature for 5 minutes. Chloroform (0.2 mL) was added and shaken for 15 seconds by hand vigorously. The mixed solution was incubated at room temperature for 3-5 minutes before centrifuged for 15 minutes at 12,000g at 4°C. The mixture was separated into a colorless upper aqueous phase, interphase, and a lower red phenol-chloroform phase and RNA remained exclusively in the upper aqueous phase. The upper aqueous phase was transferred gently into a new tube carefully. The 100% isopropanol (0.5 mL) was added to the aqueous phase and incubated after mixing for 10 minutes at room temperature. The mixture was centrifuged at 12,000g 4[for 10 minutes, and the RNA formed a gel-like pellet at the tube bottom. The supernatant was discarded, and the RNA pellet was washed twice by 75% ethanol in RNase-free water. The RNA was dissolved in 30-50 μl RNase-free water and the consistency was analyzed by NanoDrop 2000 equipment (Thermo Fisher).

### Reverse Transcription PCR

Reverse transcription PCR (RT-PCR) was performed with 500 ng RNA per reaction by TaKaRa PrimeScriptTM RT Master Mix (Perfect Real Time) Kit. Briefly, designated template solution reagent 500 ng RNA and 2 μl of PrimeScript RT Master Mix reagent were added into the 250 μl PCR tube. The RNase-Free H_2_O was added up to 10 μl for each reaction. The reagents were mixed gently, and the reverse transcription reaction was performed according to the instruction. The product was then diluted 20 times (190 μl of distilled water was added to each tube) before use.

### Quantitative Real-time PCR (QRT-PCR)

QRT-PCR was performed by the TaKaRa TB Green kit and ABI QuantStudio 12K Flex System in the 384-well plates. The primer sequences for miRNAs and genes were shown in Supplementary Table 1. The Data was analyzed via the comparison Ct (2-_ΔΔ_Ct) method and showed a fold change compared with U6 snRNA (for miRNAs) or GAPDH (for genes). Each sample was analyzed in triplicate times.

### Protein Extraction

To evaluate the protein expression, the cells were washed with 2 ml PBS three times before lysed with 60-100 μl RIPA buffer containing protease inhibitors PMSF (Thermo) on ice for 15 minutes. The cells were carefully scraped to collect all the samples into the tubes. The samples were centrifuged for 15 minutes at 12,000g and 4°C, and the supernatant was moved into new tubes. 25 μl of 5 × loading buffer was added to the protein solution and mixed well. The mixed protein samples were heated for 5 minutes at 95°C then cooled down to room temperature and stored at −80°C for the following western blotting.

### Western Blotting

The protein samples were loaded into the Tris-glycine gels (5% concentrated gel and 10% separate gel) and run in the running buffer at a constant 100V, until the bromophenol blue reaches the gel bottom. The proteins were transferred to a nitrocellulose membrane in the transfer buffer at a constant 270mA for 100 minutes. The membranes were blocked for 2 hours and incubated with primary antibody solution at 4□ overnight. The transfer membranes were washed by TBST three times and incubated with HRP-conjugated secondary antibody diluted in blocking buffer for one hour at room temperature. The transfer membranes were washed by TBST three times, and then exposed to HRP substrate (ECL solutions A and B in a 40:1 ratio) for 5-10 minutes and showed by Hyper film ECL.

### Co-immunoprecipitation and Immunoblotting

Cultured cells were lysed with cold IP-lysis buffer (20 mM Tris-HCl [pH 7.4], 0.15M NaCl, 1 mM EDTA, 1 mM EGTA, 1% Triton X-100 with protease inhibitors cocktail (Thermo) and phosphatase inhibitors (10 mM NaF, 1 mM β-glycerophosphate, 1 mM Na3VO4, 1 mM PMSF) for 30 mins and centrifuged at 13000 rounds per minute (rpm) at 4° C for 30 mins. The whole-cell lysates supernatant was transferred into new tubes. The concentration of proteins was detected by the BCA method (Thermo). The protein incubated with Anti-β-catenin or Anti-p-PKC-ζovernight at 4° C. Immunoprecipitates were washed five times with IP-lysis buffer, dissolved in sample loading buffer, and proceeded to heat denaturing and subsequently subjected to standard western immunoblot analysis.

### Chromatin immunoprecipitation

Primary chondrocytes serum-starved for 16 hours were treated with or without Wnt3a (80 ng/ml) for 1 hour and exposed to 1% formaldehyde for 10 minutes. Reactions were terminated with 0.125 M glycine. Cells were lysed in lysis buffer (50 mmol/L Tris-HCl, pH 8.0, 10 mmol/L EDTA, 1% SDS) for 1 hour and subsequently sonicated 10 times on the ice. Lysates were incubated with binding buffer (1.1% Triton-X 100, 0.1% SDS, 16.7 mmol/L Tris-HCl, 167 mmol/L NaCl, pH=8.1) with a β-catenin antibody overnight at 4°C, followed by capture with protein A-Sepharose (GE Healthcare Life Sciences) for 1 hour. Samples were washed with binding buffer and resuspended in 100μl TE before immunoblot and following QRT-PCR analyses.

### Surgically Induced OA Model

The destabilization of the medial meniscus (DMM) model is a frequently used chronic model with high clinical relevance. DMM surgery will be performed on the right knee and sham surgery on the left knee under anesthesia using Ketamine-Xylazine. A minor incision (0.5 cm) will be made on the medial parapatellar side of the right knee joint, followed by an incision of the medial capsule. Then, the medial meniscotibial ligament (MMTL) will be transected, allowing the medial meniscus to be displaced medially in knee flexion and extension positions. Finally, the joint capsule and skin will be sutured separately. As an internal control, sham surgery will be performed on the left knee joint using the same approach without MMTL transection.

### Frozen Sections

The 10% EDTA decalcified samples were rinsed thoroughly with water to eliminate the EDTA in the samples. The samples were then gradually dehydrated overnight in 10%, 20%, 30% sucrose solution. The dehydrated samples were then submerged in a solution contained an equal 30% sucrose and Optimal Cutting Temperature compound (O.C.T. Sakura 4583, Japan) overnight at 4[for preparing frozen sections. The knee joints medial compartment loading area was sectioned at 10-μm-thick sagittal-oriented in −18° C freezing microtomes (Cryostat NX70, Thermo Fisher Scientific, MA, USA) for further histological analysis.

### Safranin O and Fast Green Staining

The sections were hydrated in distilled water and stained by Weigert’s Iron Hematoxylin for 5 minutes, then washed gently in distilled water, and differentiated in 1% Acid-Alcohol for 2 seconds. The sections were rinsed gently in distilled water three times and stained by 0.2% Fast Green for 3 minutes, then rinsed quickly with 1% acetic acid solution for 10 seconds and stained in 0.1% safranin O solution for 15 minutes. The sections were dehydrated and cleared with 80%, 90%, 95% ethyl alcohol, and absolute ethyl alcohol, and xylene twice for 2 minutes each time, then mounted in the resinous medium.

### OA Evaluation under OARSI Grade Guideline

The OA evaluation was based on a well-established OARSI (Osteoarthritis Research Society International) guideline which contained both photograph reference and text instruction. with the Safranin O and Fast Green stained sections. The OA evaluation was performed by two double-blinded colleagues.

### Immunofluorescence

The sections were washed gently by PBS, permeabilized by 0.2% Triton-X for 10 minutes, washed gently in PBS, and blocked at room temperature for 2 hours by block solution (1% Bovine Serum Albumin). The sections were then incubated with primary antibodies in block solution overnight at 4° C. The sections were washed gently with 0.2% Triton-X for 1 minute to reduce the background and washed by PBS. The sections were incubated with secondary antibody for 1 hour at room temperature and protect from light, then washed gently with 0.2% Triton-X for 1 minute to reduce the background and washed by PBS. The sections were stained with DAPI solution (1:2000) for 5 minutes, then washed in PBS three times, and finally mounted (F4680, Sigma-Aldrich. St. Louis, MO).

### Sample Size Estimation for in vivo Experiment

The sample size estimation was based on the osteoarthritis research by the same animal model and power analysis of the results. The histological analysis under OARSI guideline was used to analyze the statistical power of this study. The statistical power of 0.80 with significance level set at 0.05 (two-tail) to detect the difference was chosen to evaluate the sample size by the G*Power v3.1 software. Therefore, the sample size equals 8 provides sufficient statistical power to detect the difference between the control and treatment groups.

### Statistical Analysis

Data were shown as the mean ± standard deviation (SD). For the comparison between the two groups, the unpaired two-tailed Student’s t-test was carried out. And for multiple group comparisons, one-way or two-way ANOVA was used with Tukey *post hoc* test. p < 0.05 was considered as statistically significant. GraphPad Prism software (Version 8.3.0) was used for statistical analysis.

## Author contribution

W.T. performed most of the experiments and analyzed the data. J.X. and Q.Q partly designed the experiments and helped to perform the animal surgery. H.C., T.H., C.C., and W.L. partly performed the in vitro experiments. Z.H., Y.C., Z.M., and D.Z. collected the clinical specimens. W.T., J.X., and J.H. were involved in manuscript preparation. J.H. provided the conceptual framework, designed the study, supervised the project, and wrote the manuscript.

## Acknowledgments (Funding)

The work was supported by the National Natural Science Foundation of China (Ref No. 82002359), Guangdong Basic and Applied Basic Research Foundation (Ref No. 2020A1515010267), China Postdoctoral Science Foundation (Ref No. 2019M663020), Grant for Key Disciplinary Project of Clinical Medicine under the Guangdong High-level University Development Program (Ref No. 002-18119101 and 002-18120303), “Dengfeng Project” for the construction of high-level hospitals in Guangdong Province-the First Affiliated Hospital of Shantou University Medical College Supporting Funding, Li Ka Shing Foundation Cross-Disciplinary Research Grant (Ref No. 2020LKSFG03A). The funding bodies had no role in the design of the study, the collection, analysis, interpretation of the data, writing the manuscript.

## Patient consent

Obtained

## Ethics approval

All animal experiments were conducted under the approval of the animal ethics committee in our institute by following the guidelines. The tissue bank for the study of degenerative joint disease was approved by the clinical research committee as well.

---

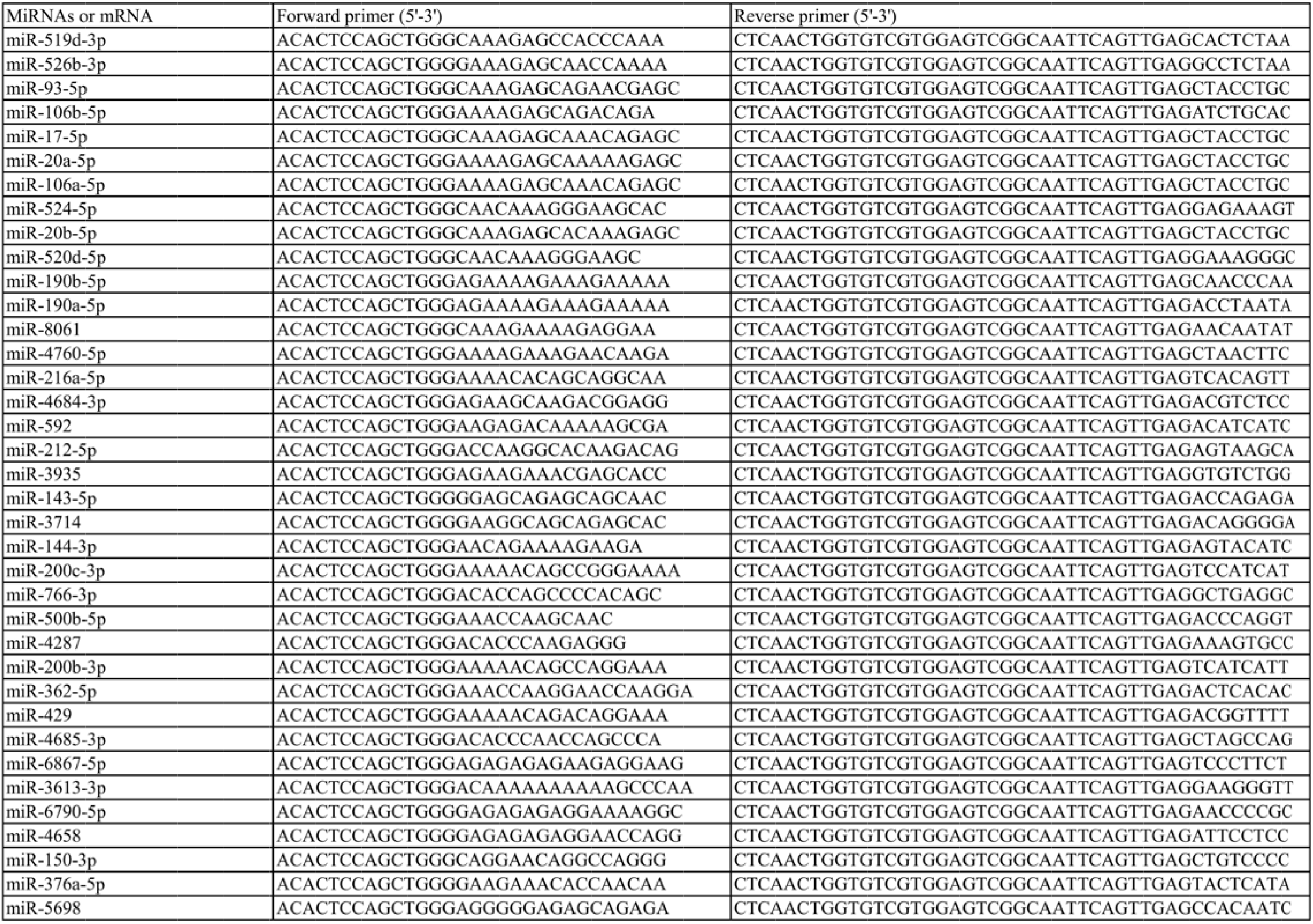
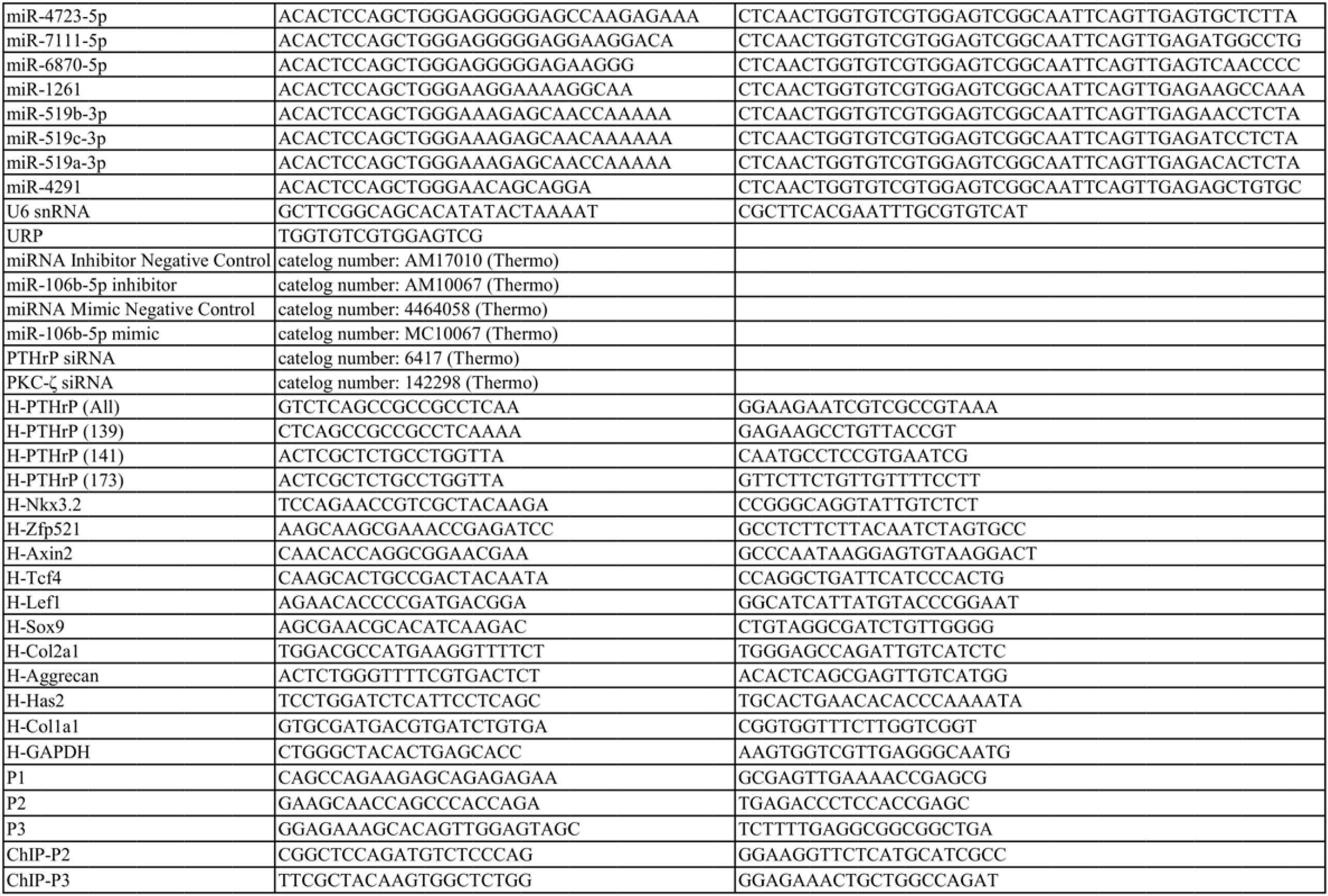

## References

1. S. L. James, D. Abate, K. H. Abate, S. M. Abay, C. Abbafati, N. Abbasi, H. Abbastabar, F. Abd-Allah, J. Abdela, A. Abdelalim, Global, regional, and national incidence, prevalence, and years lived with disability for 354 diseases and injuries for 195 countries and territories, 1990–2017: a systematic analysis for the Global Burden of Disease Study 2017. The Lancet 392, 1789–1858 (2018).

2. R. Raouf, S. Lolignier, J. E. Sexton, Q. Millet, S. Santana-Varela, A. Biller, A. M. Fuller, V. Pereira, J. S. Choudhary, M. O. Collins, Inhibition of somatosensory mechanotransduction by annexin A6. Science signaling 11, (2018).

3. E. M. Roos, N. K. Arden, Strategies for the prevention of knee osteoarthritis. Nature Reviews Rheumatology 12, 92 (2016).

4. W. Tong, Y. Geng, Y. Huang, Y. Shi, S. Xiang, N. Zhang, L. Qin, Q. Shi, Q. Chen, K. Dai, In vivo identification and induction of articular cartilage stem cells by inhibiting NF[κB signaling in osteoarthritis. Stem Cells 33, 3125–3137 (2015).

5. Q. Wang, C. M. Lepus, H. Raghu, L. L. Reber, M. M. Tsai, H. H. Wong, E. von Kaeppler, N. Lingampalli, M. S. Bloom, N. Hu, IgE-mediated mast cell activation promotes inflammation and cartilage destruction in osteoarthritis. Elife 8, e39905 (2019).

6. H. Clevers, R. Nusse, Wnt/β-catenin signaling and disease. Cell 149, 1192–1205 (2012).

7. J. Yang, M. Kitami, H. Pan, M. T. Nakamura, H. Zhang, F. Liu, L. Zhu, Y. Komatsu, Y. Mishina, Augmented BMP signaling commits cranial neural crest cells to a chondrogenic fate by suppressing autophagic β-catenin degradation. Science Signaling 14, (2021).

8. S. Park, L. Wu, J. Tu, W. Yu, Y. Toh, K. S. Carmon, Q. J. Liu, Unlike LGR4, LGR5 potentiates Wnt–β-catenin signaling without sequestering E3 ligases. Science Signaling 13, (2020).

9. E. Canalis, Wnt signalling in osteoporosis: mechanisms and novel therapeutic approaches. Nature Reviews Endocrinology 9, 575–583 (2013).

10. Y. Usami, A. T. Gunawardena, M. Iwamoto, M. Enomoto-Iwamoto, Wnt signaling in cartilage development and diseases: lessons from animal studies. Laboratory investigation 96, 186–196 (2016).

11. Z.-c. Ding, Y.-k. Lin, Y.-k. Gan, T.-t. Tang, Molecular pathogenesis of fracture nonunion. Journal of orthopaedic translation 14, 45–56 (2018).

12. R. J. Lories, M. Corr, N. E. Lane, To Wnt or not to Wnt: the bone and joint health dilemma. Nature Reviews Rheumatology 9, 328 (2013).

13. M. Zhu, D. Tang, Q. Wu, S. Hao, M. Chen, C. Xie, R. N. Rosier, R. J. O’Keefe, M. Zuscik, D. Chen, Activation of β[catenin signaling in articular chondrocytes leads to osteoarthritis[like phenotype in adult β[catenin conditional activation mice. Journal of Bone and Mineral Research 24, 12–21 (2009).

14. T. Yuasa, N. Kondo, R. Yasuhara, K. Shimono, S. Mackem, M. Pacifici, M. Iwamoto, M. Enomoto-Iwamoto, Transient activation of Wnt/β-catenin signaling induces abnormal growth plate closure and articular cartilage thickening in postnatal mice. The American journal of pathology 175, 1993–2003 (2009).

15. M. Zhu, M. Chen, M. Zuscik, Q. Wu, Y. J. Wang, R. N. Rosier, R. J. O’Keefe, D. Chen, Inhibition of β[catenin signaling in articular chondrocytes results in articular cartilage destruction. Arthritis & Rheumatism: Official Journal of the American College of Rheumatology 58, 2053–2064 (2008).

16. G. Nalesso, B. L. Thomas, J. C. Sherwood, J. Yu, O. Addimanda, S. E. Eldridge, A.-S. Thorup, L. Dale, G. Schett, J. Zwerina, WNT16 antagonises excessive canonical WNT activation and protects cartilage in osteoarthritis. Annals of the rheumatic diseases 76, 218–226 (2017).

17. A. J. Mikels, R. Nusse, Purified Wnt5a protein activates or inhibits β-catenin–TCF signaling depending on receptor context. PLoS Biol 4, e115 (2006).

18. B. Shu, Y. Zhao, S. Zhao, H. Pan, R. Xie, D. Yi, K. Lu, J. Yang, C. Xue, J. Huang, Inhibition of Axin1 in osteoblast precursor cells leads to defects in postnatal bone growth through suppressing osteoclast formation. Bone research 8, 1–8 (2020).

19. E. Schipani, S. Provot, PTHrP, PTH, and the PTH/PTHrP receptor in endochondral bone development. Birth Defects Research Part C: Embryo Today: Reviews 69, 352–362 (2003).

20. H. M. Kronenberg, PTHrP and skeletal development. Annals of the New York Academy of Sciences 1068, 1–13 (2006).

21. W. Huang, U.-i. Chung, H. M. Kronenberg, B. de Crombrugghe, The chondrogenic transcription factor Sox9 is a target of signaling by the parathyroid hormone-related peptide in the growth plate of endochondral bones. Proceedings of the National Academy of Sciences 98, 160–165 (2001).

22. W. Tong, Y. Zeng, D. H. K. Chow, W. Yeung, J. Xu, Y. Deng, S. Chen, H. Zhao, X. Zhang, K. K. Ho, Wnt16 attenuates osteoarthritis progression through a PCP/JNK-mTORC1-PTHrP cascade. Annals of the rheumatic diseases 78, 551–561 (2019).

23. C. Macica, G. Liang, A. Nasiri, A. E. Broadus, Genetic evidence of the regulatory role of parathyroid hormone–related protein in articular chondrocyte maintenance in an experimental mouse model. Arthritis & Rheumatism 63, 3333–3343 (2011).

24. X. Chen, C. M. Macica, B. E. Dreyer, V. E. Hammond, J. R. Hens, W. M. Philbrick, A. E. Broadus, Initial characterization of PTH[related protein gene[driven lacZ expression in the mouse. Journal of Bone and Mineral Research 21, 113–123 (2006).

25. X. Chen, C. M. Macica, A. Nasiri, A. E. Broadus, Regulation of articular chondrocyte proliferation and differentiation by Indian hedgehog and parathyroid hormone–related protein in mice. Arthritis & Rheumatism: Official Journal of the American College of Rheumatology 58, 3788–3797 (2008).

26. C. Becher, T. Szuwart, P. Ronstedt, S. Ostermeier, A. Skwara, S. Fuchs-Winkelmann, C. O. Tibesku, Decrease in the expression of the type 1 PTH/PTHrP receptor (PTH1R) on chondrocytes in animals with osteoarthritis. Journal of orthopaedic surgery and research 5, 1–6 (2010).

27. K. Gelse, A. Ekici, F. Cipa, B. Swoboda, H. Carl, A. Olk, F. Hennig, P. Klinger, Molecular differentiation between osteophytic and articular cartilage–clues for a transient and permanent chondrocyte phenotype. Osteoarthritis and cartilage 20, 162–171 (2012).

28. D. Wang, J. M. Taboas, R. S. Tuan, PTHrP overexpression partially inhibits a mechanical strain-induced arthritic phenotype in chondrocytes. Osteoarthritis and Cartilage 19, 213–221 (2011).

29. J. K. Chang, L. H. Chang, S. H. Hung, S. C. Wu, H. Y. Lee, Y. S. Lin, C. H. Chen, Y. C. Fu, G. J. Wang, M. L. Ho, Parathyroid hormone 1–34 inhibits terminal differentiation of human articular chondrocytes and osteoarthritis progression in rats. Arthritis & Rheumatism: Official Journal of the American College of Rheumatology 60, 3049–3060 (2009).

30. W. Zhang, J. Chen, S. Zhang, H. W. Ouyang, Inhibitory function of parathyroid hormone-related protein on chondrocyte hypertrophy: the implication for articular cartilage repair. Arthritis research & therapy 14, 1–10 (2012).

31. X. Guo, K. K. Mak, M. M. Taketo, Y. Yang, The Wnt/β-catenin pathway interacts differentially with PTHrP signaling to control chondrocyte hypertrophy and final maturation. PloS one 4, e6067 (2009).

32. M. Hiremath, P. Dann, J. Fischer, D. Butterworth, K. Boras-Granic, J. Hens, J. Van Houten, W. Shi, J. Wysolmerski, Parathyroid hormone-related protein activates Wnt signaling to specify the embryonic mammary mesenchyme. Development 139, 4239–4249 (2012).

33. R. W. Johnson, A. R. Merkel, J. M. Page, N. S. Ruppender, S. A. Guelcher, J. A. Sterling, Wnt signaling induces gene expression of factors associated with bone destruction in lung and breast cancer. Clinical & experimental metastasis 31, 945–959 (2014).

34. K. K. Mak, H. M. Kronenberg, P.-T. Chuang, S. Mackem, Y. Yang, Indian hedgehog signals independently of PTHrP to promote chondrocyte hypertrophy. Development 135, 1947–1956 (2008).

35. T. Kobayashi, U.-i. Chung, E. Schipani, M. Starbuck, G. Karsenty, T. Katagiri, D. L. Goad, B. Lanske, H. M. Kronenberg, PTHrP and Indian hedgehog control differentiation of growth plate chondrocytes at multiple steps. Development 129, 2977–2986 (2002).

36. J. Jiang, N. Leong, J. Mung, C. Hidaka, H. Lu, Interaction between zonal populations of articular chondrocytes suppresses chondrocyte mineralization and this process is mediated by PTHrP. Osteoarthritis and Cartilage 16, 70–82 (2008).

37. R. W. Johnson, M. P. Nguyen, S. S. Padalecki, B. G. Grubbs, A. R. Merkel, B. O. Oyajobi, L. M. Matrisian, G. R. Mundy, J. A. Sterling, TGF-β promotion of Gli2-induced expression of parathyroid hormone-related protein, an important osteolytic factor in bone metastasis, is independent of canonical Hedgehog signaling. Cancer research 71, 822–831 (2011).

38. C. Zhang, X. Wei, C. Chen, K. Cao, Y. Li, Q. Jiao, J. Ding, J. Zhou, B. C. Fleming, Q. Chen, Indian hedgehog in synovial fluid is a novel marker for early cartilage lesions in human knee joint. International journal of molecular sciences 15, 7250–7265 (2014).

39. F. Wei, J. Zhou, X. Wei, J. Zhang, B. C. Fleming, R. Terek, M. Pei, Q. Chen, T. Liu, L. Wei, Activation of Indian hedgehog promotes chondrocyte hypertrophy and upregulation of MMP-13 in human osteoarthritic cartilage. Osteoarthritis and cartilage 20, 755–763 (2012).

40. Y. Shi, G. He, W.-C. Lee, J. A. McKenzie, M. J. Silva, F. Long, Gli1 identifies osteogenic progenitors for bone formation and fracture repair. Nature communications 8, 2043 (2017).

41. L. N. Reynard, M. J. Barter, Osteoarthritis year in review 2019: genetics, genomics and epigenetics. Osteoarthritis and Cartilage 28, 275–284 (2020).

42. R. Garg, L. G. Benedetti, M. B. Abera, H. Wang, M. Abba, M. G. Kazanietz, Protein kinase C and cancer: what we know and what we do not. Oncogene 33, 5225–5237 (2014).

43. C. Matta, A. Mobasheri, Regulation of chondrogenesis by protein kinase C: emerging new roles in calcium signalling. Cellular signalling 26, 979–1000 (2014).

44. R. Kc, X. Li, J. S. Kroin, Z. Liu, D. Chen, G. Xiao, B. Levine, J. Li, J. L. Hamilton, A. J. van Wijnen, PKCδ null mutations in a mouse model of osteoarthritis alter osteoarthritic pain independently of joint pathology by augmenting NGF/TrkA-induced axonal outgrowth. Annals of the rheumatic diseases 75, 2133–2141 (2016).

45. X. Martineau, É. Abed, J. Martel-Pelletier, J.-P. Pelletier, D. Lajeunesse, Alteration of Wnt5a expression and of the non-canonical Wnt/PCP and Wnt/PKC-Ca2+ pathways in human osteoarthritis osteoblasts. PloS one 12, e0180711 (2017).

46. C. Matta, A. Mobasheri, P. Gergely, R. Zákány, Ser/Thr-phosphoprotein phosphatases in chondrogenesis: neglected components of a two-player game. Cellular signalling 26, 2175–2185 (2014).

47. Y. Hong, aPKC: The kinase that phosphorylates cell polarity. F1000Research 7, (2018).

48. A. Suzuki, S. Ohno, The PAR-aPKC system: lessons in polarity. Journal of cell science 119, 979–987 (2006).

49. A. X. Wu-Zhang, A. C. Newton, Protein kinase C pharmacology: refining the toolbox. Biochemical Journal 452, 195–209 (2013).

